# SPECtre: a spectral coherence-based classifier of actively translated transcripts from ribosome profiling sequence data

**DOI:** 10.1101/034777

**Authors:** Sang Y. Chun, Caitlin M. Rodriguez, Peter K. Todd, Ryan E. Mills

**Affiliations:** Department of Computational Medicine and Bioinformatics, University of Michigan, Ann Arbor, MI, 48109, USA; Department of Neurology, University of Michigan, Ann Arbor, MI, 48109, USA; Veterans Affairs Medical Center, Ann Arbor, MI, 48105; Department of Human Genetics, University of Michigan, Ann Arbor, MI, 48109, USA

## Abstract

**Summary:** Active protein translation can be assessed and measured using ribosome profiling sequencing strategies. Existing analytical approaches applied to this technology make use of sequence fragment length or frame occupancy to differentiate between active translation and background noise, however they do not consider additional characteristics inherent to the technology which limits their overall accuracy. Here, we present an analytical tool that models the overall tri-nucleotide periodicity of ribosomal occupancy using a classifier based on spectral coherence. Our software, SPECtre, examines the relationship of normalized ribosome profiling read coverage over a rolling series of windows along a transcript against an idealized reference signal. A comparison of SPECtre against current methods on existing and new data shows a marked improvement in accuracy for detecting active translation and exhibits overall high sensitivity at a low false discovery rate.

**Availability and Implementation:** SPECtre source code is available for download at https://github.com/mills-lab/spectre.

## 1 Introduction

Ribosome profiling is a next-generation sequencing strategy that enriches for ribosome-protected mRNA footprints indicative of active protein translation (Ingolia, 2009). Fragments of mRNA bound by ribosomal complexes are selected for by enzymatic digestion, isolated using a sucrose cushion or gradient, released from their occupying ribosome, size-selected by gel electrophoresis, and then sequenced. Thus, sequencing and analysis of ribosome-protected fragments of mRNA enables profiling of the translational content of a sample on a transcriptome-wide level.

Various algorithms have been developed to differentiate protein-coding and non-coding transcripts in ribosome profiling sequence data using fragment length distribution differences (Ingolia, 2014) and read frame alignment enrichment (Bazzini, 2014). However, classification based on extreme outlier analysis of fragment length organization similarity score (FLOSS) differences is agnostic to the ribosome-protected fragment abundance over a transcript. Furthermore, classification based on read frame alignment enrichment (ORFscore) is optimized for canonical open reading frame usage only. In addition, neither of the algorithms described above are available as standalone packages and must be implemented by the user.

Here we introduce SPECtre, a spectral coherence-based classification algorithm to identify regions of active translation using aligned ribosome profiling sequence reads (Figure 1A). SPECtre leverages a key feature of ribosome profiling where sequence reads aligned to a reference transcriptome will track the tri-nucleotide periodicity characteristic of ` transcripts as they are translated by ribosomes. Options to change the step size between windows, the size of windows analyzed, false discovery rate and abundance cutoffs to differentiate translated versus non-translated distributions are provided to the end-user to customize. Implementations of FLOSS and ORFscore are included for comparative purposes.

**Fig. 1.**
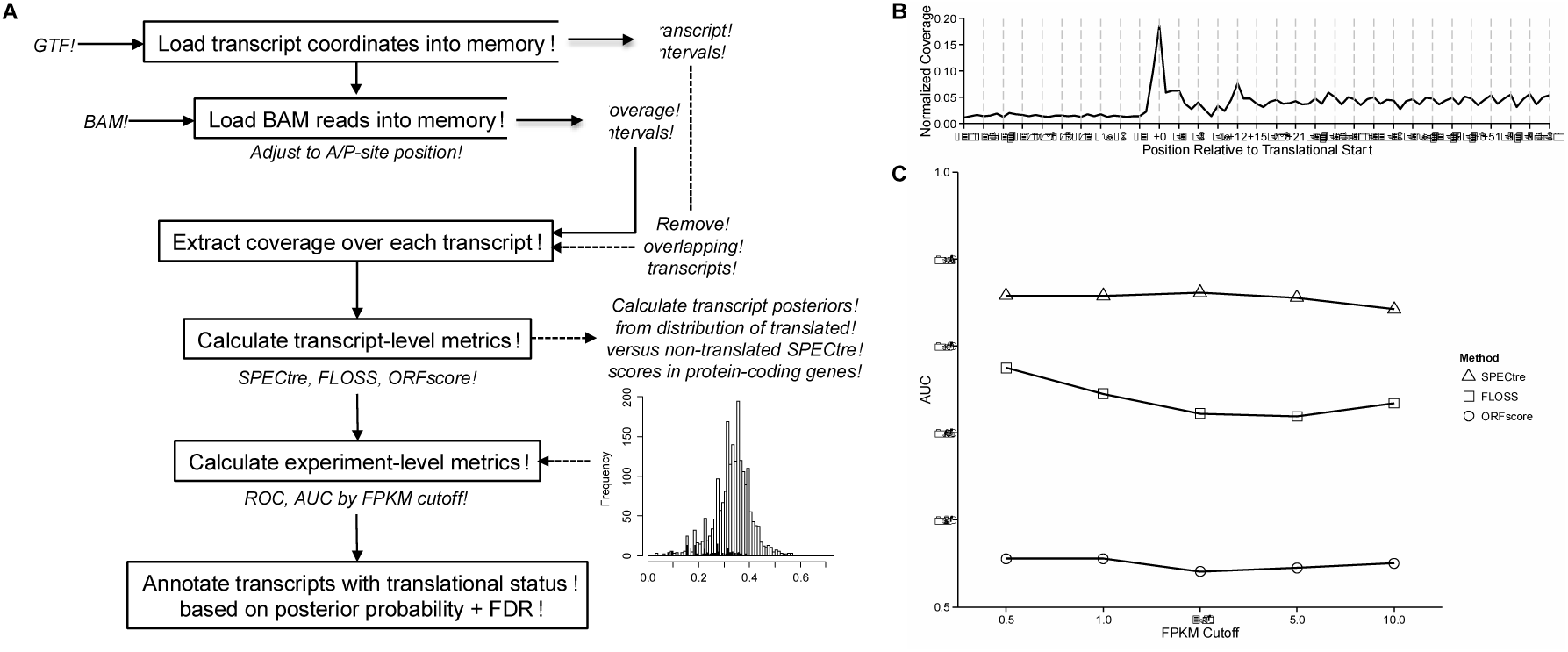
(A) Overview of the SPECtre analytical pipeline. (B) Normalized ribosome profiling read coverage aggregated over all transcripts in human SH-SY5Y neuroblastoma cells indicating regions of active translation as characterized by the tri-nucleotide periodic signal. (C) Comparative analysis of classifier method performance in human SH-SY5Y neuroblastoma cells at different minimum FPKM cutoffs. Points shown denote the AUC for ROC classification using SPECtre (triangle), FLOSS (square), and ORFscore (circle).

## 2 Methods

In contrast to non-coding transcripts, ribosome profiling sequence reads aligned to coding transcripts are characterized by a tri-nucleotide periodic signal as ribosome-bound mRNA is translated into protein in a codon-dependent manner (Figure 1B). Thus coding transcripts may be differentiated from non-coding transcripts by the presence or absence of a strong tri-nucleotide periodic signal. To measure the strength of this tri-nucleotide signal, we calculate the spectral coherence (Bendat, 1986) over sliding N nucleotide windows over a transcript. Spectral coherence is a measurement of the power relationship between two signals over the frequency domain, such that two signals with shared frequencies will have high coherence, whereas two unrelated signals will be of low coherence. The SPECtre score, based on the spectral coherence over overlapping windows across a transcript, is calculated for each transcript from a user-provided transcript annotation database. Distributions of these scores are generated using a user-defined FPKM cutoff to delineate transcripts under active translation versus those that are not, and these are then used to derive a minimum SPECtre score threshold for active translation given a customizable false discovery rate.

## 3 Results

We assessed the sensitivity and specificity of each classification algorithm in previously published ribosome profiling data for mouse (Ingolia, 2014) and zebrafish (Bazzini, 2014). We also used data generated in house from a human SH-SY5Y neuroblastoma cell line (See Supplemental Materials, Supplemental Table 1). Ribosome profiling sequence reads from each set were aligned to their respective genome and transcriptome reference. Overlapping and neighboring annotated protein-coding and non-coding transcripts were removed from the analysis using methods described previously (Ingolia, 2014). The FLOSS, ORFscore and SPECtre metrics were calculated for each transcript in the aforementioned sanitized transcript set using the default parameters; the median SPECtre score was calculated over 30 nucleotide overlapping windows, with 3 nucleotides stepped to the next window in order to capture the canonical reading frame. Next, we re-labeled annotated protein-coding transcripts as either ‘translated’ or ‘not translated’ based on their abundance of ribosome-protected fragments over a series of minimum FPKM (Trapnell, 2010) cutoffs. In this manner, we are able to assess the ability of each approach to identify transcripts with signatures of active translation in the interrogated cell type.

We performed receiver operating characteristic (ROC) analyses and calculated the area under the curve (AUC) for each of the three classification algorithms over variable minimum FPKM cutoffs. SPECtre classification outperforms both FLOSS and ORFscore to identify actively translated protein-coding transcripts in a ribosome profiling library generated from human neuroblastoma cells (Figure 1C), and remains robust in a mouse embryonic stem cell ribosome profiling library (Supplemental Figure 1A). In addition, SPECtre exhibits a marked improvement in accuracy in a meta-analysis of ribosome profiling libraries derived from zebrafish embryos (Supplemental Figure 1B). In summary, SPECtre is a flexible, command-line-driven analytical package that identifies regions of active translation in ribosome profiling sequence data, and is robust across ribosome profiling libraries derived from multiple organisms and cell types.

In summary, SPECtre is a flexible, command-line-driven analytical package that identifies regions of active translation in ribosome profiling sequence data, and is robust across ribosome profiling libraries derived from multiple organisms and cell types.

